# Breaking the Link: Neural Dissociation of Attention and Working Memory through Inhibitory Control

**DOI:** 10.1101/2025.04.17.649277

**Authors:** Yueyao Liu, Yingtao Fu, Enze Tang, Hang Wu, Junrong Han, Musi Xie, Yihui Zhang, Jinhao Huang, Hanjun Liu, Hui Chen, Pengmin Qin

## Abstract

Attention and working memory (WM) encoding have traditionally been considered inseparable processes with shared neural mechanisms. Here, using an innovative experimental design and a multimodal approach, we provide the first direct neural evidence that attention and WM encoding are dissociable. Functional MRI identifies the supramarginal gyrus (SMG) as the key region enabling this dissociation, while dynamic causal modeling reveals the neural circuitry through which the SMG exerts inhibitory control over attentional representations, regulating their integration into WM. Furthermore, neuromodulation via transcranial direct current stimulation (tDCS) demonstrates that enhancing SMG activity strengthens this inhibitory control, providing causal evidence for the dissociation mechanism. A second tDCS experiment with varied stimuli confirms the generalizability of this mechanism and reinforces the robustness of our results. These findings challenge the long-standing view that attention and WM encoding form a continuous process, demonstrating instead that they constitute two dissociable neural processes of information selection.

## Introduction

Attention and working memory (WM) are fundamental cognitive functions that support goal-directed behavior^1^. Attention enables individuals to focus on relevant information in complex environments, while WM temporarily stores selected inputs for cognitive processing. Understanding their interaction is critical for uncovering how the brain prioritizes, processes, and retains information. Traditionally, attention and WM have been considered tightly linked in their neural underpinnings. Neuroimaging and electrophysiological studies in humans and macaques suggest that attentional selection and WM processes share a common neural basis, primarily in the prefrontal and parietal cortices^2–6^.

However, recent evidence suggests that attention and WM may rely on distinct neural mechanisms. Mendoza-Halliday et al. (2024) trained macaques to either maintain features in WM or use them to guide attentional deployment, identifying separate neuronal substrates in the posterior lateral prefrontal cortex for WM storage and feature-based attention^7^. This finding highlights the dissociation between maintaining WM representations and using them to guide attentional allocation. However, traditional models emphasizing the close interdependence between attention and WM have primarily focused on another aspect—the relationship between attentional selection and WM encoding. From the traditional perspective, attention has long been regarded as the gatekeeper of WM, with the assumption that attended information is automatically encoded into WM^3,8–10^. In conventional experimental designs, attentional selection and WM encoding have been difficult to dissociate, as attended information is typically required to be encoded into WM^5,6,10^.

Here, we introduce a novel design to address this limitation, enabling a direct dissociation between attentional selection and WM encoding within a single behavioral task^11,12^. This method provides a unique opportunity to examine the neural mechanisms underlying their dissociation. Understanding this neural separation is crucial, as it challenges the long-standing assumption that attention and WM constitute a continuous process rather than two distinct stages of information selection.

Based on this new design, we employ a multimodal approach integrating neuroimaging, computational modeling, and neurostimulation. Using functional MRI (fMRI), we identify brain regions involved in regulating the transition of attended representations into WM, distinguishing those that facilitate this transfer from those that suppress unnecessary encoding. To further explore the dynamic neural interactions governing this process, we develop computational models of neural circuits to examine how different brain regions coordinate WM integration for attended representations. Finally, we apply transcranial direct current stimulation (tDCS) to modulate activity in key regions identified through fMRI and computational modeling, providing causal evidence for how neural activity influences the gating of representations from attention to WM. To ensure the robustness of our findings, we replicate the tDCS experiments across diverse stimulus types. Uncovering this dissociable neural basis redefines our understanding of how the brain prioritizes and organizes information to support cognition.

## Results

### Attended information can be blocked from working memory

To investigate the neural mechanisms underlying the dissociation between attentional selection and WM encoding, we conducted an fMRI experiment using a novel design modified from the attribute amnesia paradigm^11^ to isolate these two processes. As illustrated in Fig. 1, participants were tested under two experimental conditions: the pre-surprise condition and the post-surprise condition. They performed the same primary task in both conditions; however, the expectation of a memory report on the attended information varied.

**Fig. 1.**
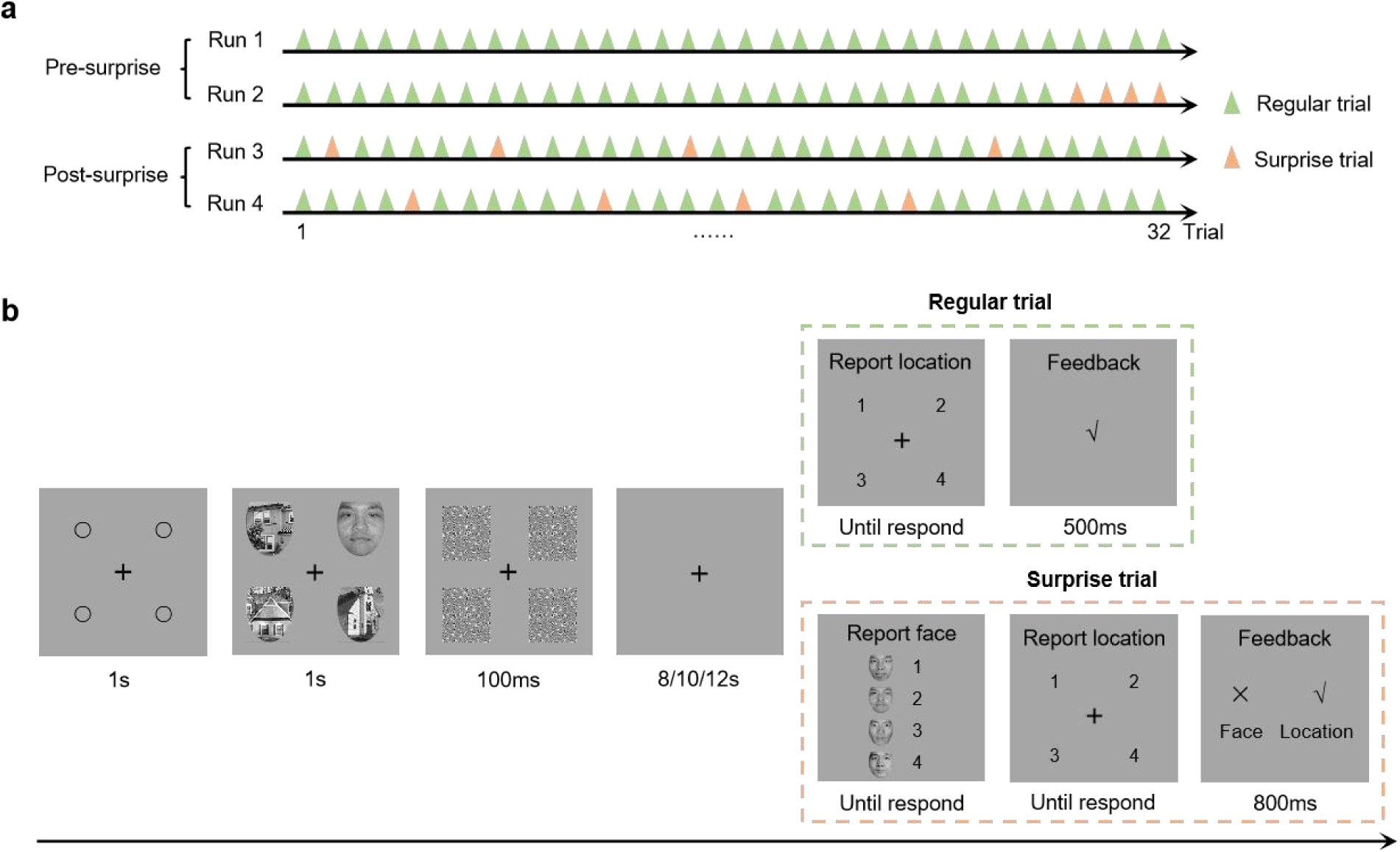
fMRI experimental design and task procedure. (**a**). The fMRI experiment comprised four runs, with 32 trials per run. The pre-surprise condition included 60 regular trials (green triangles), while the post-surprise condition included 56 regular trials. Surprise trials (orange triangles) served as catch trials to monitor face processing and maintain the expectation of reporting faces. (**b**). In regular trials, participants are instructed to report the location of the face. In surprise trials, participants are instructed to select the target faces and then report the location. The instruction for the location-reporting task is “Press a corresponding number to indicate the location of the face.” The instruction for the face-reporting task is “This is a surprise memory test! Press a corresponding number to indicate which one is the face you just saw.” The copyright for the Chinese Facial Affective Picture System used in the study has been duly purchased^61^.

In the pre-surprise condition, participants completed a face location-reporting task (regular trials) requiring visual searches within a stimulus array. Face images served as target stimuli and were fully attended and processed within a few hundred milliseconds. However, participants were explicitly instructed to remember only the location of the face and to report the corresponding location number. Consequently, the face images represented attended-without-memory information—fully attended but not necessarily stored for memory report.

After completing 60 regular trials, participants encountered four surprise trials. In the first trial, they were unexpectedly asked to select the face image they had seen moments earlier. This assessed whether participants, lacking the expectation to report the face, had refrained from storing it in working memory. The subsequent three trials served as controls, examining whether participants, after developing a reporting expectation, began encoding the face images into working memory. These catch trials disrupted participants’ prior expectations and redefined the memory relevance of the face stimuli.

In the post-surprise condition, participants continued the same face location-reporting task as in the pre-surprise condition. However, following the surprise test, participants expected that they would be required to report the target face. This expectation was reinforced by the random insertion of surprise trials. While the primary task of location-reporting remained unchanged, participants actively stored the target faces in working memory, with the face images thus becoming attended-with-memory information.

According to the behavioral results, participants maintained high accuracy in the location task, with 97.64% accuracy. This indicates that they accurately reported the target location and remained fully engaged in the experiment. As shown in Table 1, the accuracy of the face-reporting task in the first surprise trial was 48.98% (24/49), significantly lower than in subsequent surprise trials (all *p* < 0.05, Chi-squared Test). These results indicate that participants struggled to report the target face when it was attended-without-memory information but showed significant improvement in subsequent trials when it became attended-with-memory information. These findings validate the effectiveness of this design in dissociating attention from WM.

**Table 1.** Behavioral results from surprise trials in the fMRI experiment Note. The accuracy of the face-reporting task in the first surprise trial in Run 2 was significantly lower than in subsequent surprise trials, with all *p* < 0.05 by Chi-squared Test.

|  | First | Second | Third | Fourth |
| --- | --- | --- | --- | --- |
| Run 2 | 48.98% | 83.67% | 81.63% | 81.63% |
| Run 3 | 77.55% | 91.84% | 69.39% | 73.47% |
| Run 4 | 73.47% | 87.76% | 87.76% | 75.51% |

### Stronger activation of the SMG in the pre-surprise condition

Whole-brain analyses using a general linear model compared blood oxygen-level-dependent (BOLD) signals between the two conditions (see Methods for details). We identified brain regions involved in regulating the transition of attended representations into WM, distinguishing those that facilitate this transfer from those that suppress unnecessary memory encoding. As shown in Fig. 2, the supramarginal gyrus (SMG) exhibited significantly greater activation in the pre-surprise condition than in the post-surprise condition, suggesting its role in inhibiting target faces from unnecessary memory encoding. Conversely, the dorsolateral prefrontal cortex (dlPFC), supplementary motor area (SMA), and ventromedial prefrontal cortex (vmPFC) showed stronger activation in the post-surprise condition, indicating their involvement in the memory storage of target faces.

**Fig. 2.**
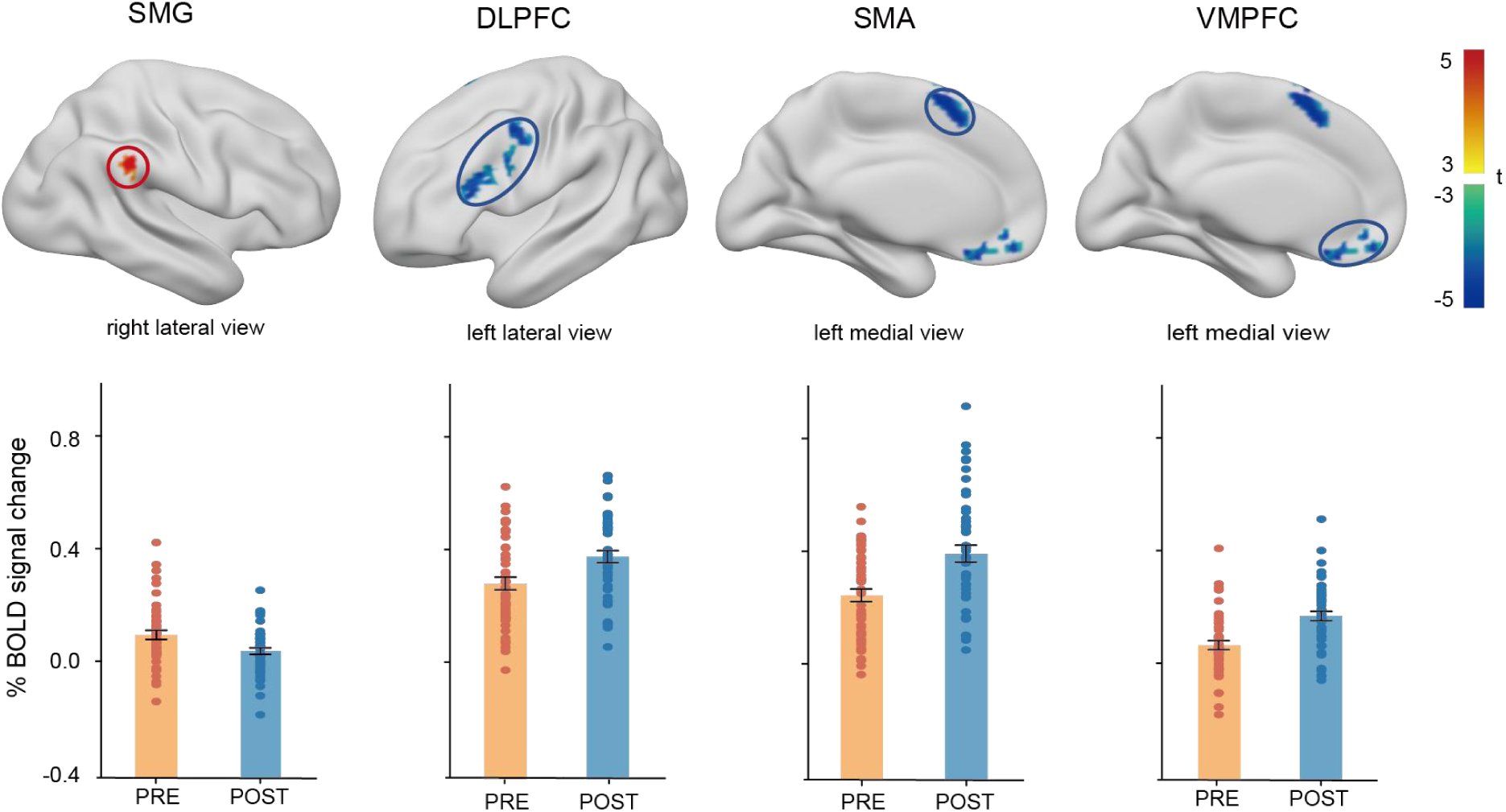
Whole-brain analyses for the fMRI experiment. A general linear model analysis was employed to compare whole-brain BOLD signals between the two conditions, with results shown for the contrast “pre-surprise > post-surprise”. A paired t-test was applied with a significance threshold of *p* < 0.01, FDR correction at the voxel level, and a cluster size of > 40 voxels. Positive activation (red) indicates stronger activation in the pre-surprise condition, whereas negative activation (blue) indicates higher activation in the post-surprise condition. Error bars indicate SEM calculated across participants and colored dots indicate data from individual participants. Abbreviations: dlPFC, dorsolateral prefrontal cortex; SMA, supplementary motor area; vmPFC, ventromedial prefrontal cortex; SMG, supramarginal gyrus.

### Neural representation of face perception when it is attended without or with memory

Based on previous studies of the automatic attentional advantages of face processing^13–17^ and the attribute amnesia effect^11,18,19^, both category and identity representations are expected to be processed during the target face-location task in both conditions. Consequently, we focused on two key perceptual processing regions as regions of interest (ROIs) to examine face perception in the brain. These ROIs were identified as the anterior temporal lobe (ATL), responsible for processing face identity^20,21^, and the fusiform face area (FFA), responsible for category representations of faces^22,23^.

Therefore, after completing the attribute amnesia experiment, participants also underwent a localizer task during the fMRI scan, specifically designed to identify the coordinate positions of the face perception regions. The localizer task employed a block design, with half of the blocks presenting face images and the other half presenting house images^24^. Participants were instructed to perform a one-back task in order to maintain focus on the stimuli. A general linear model analyses was applied to identify voxels with higher activation for faces. The localization criterion for the ATL involved the activated voxel cluster in the anterior part of the temporal lobe, while for the FFA, it was the activated voxel cluster in the middle fusiform gyrus^20^. Following these criteria, we successfully localized both the bilateral ATL and bilateral FFA, as depicted in Fig. 3.

**Fig. 3.**
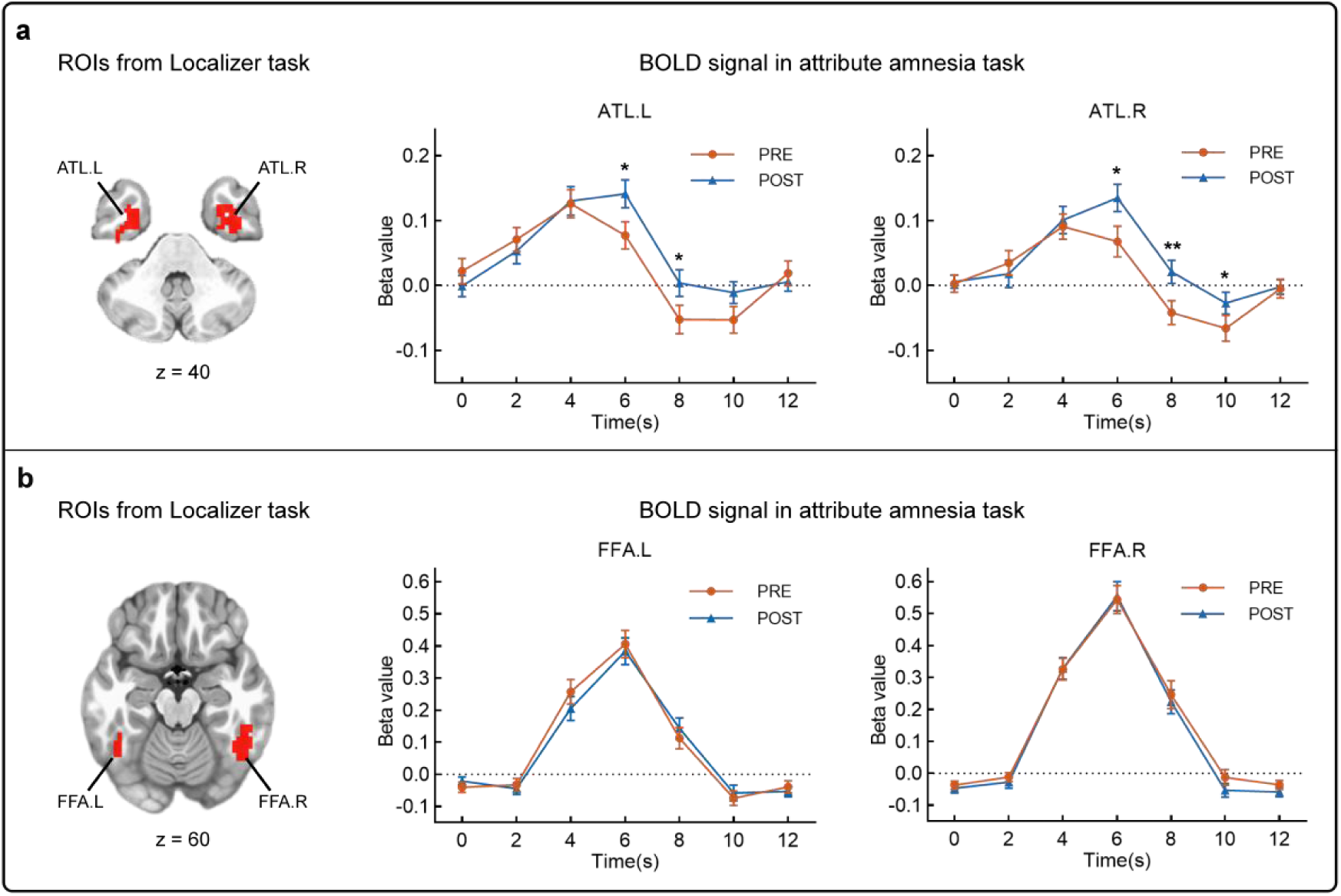
ROI analysis for face perception. The ROIs for bilateral ATL (a) and FFA (b) are shown on the left side of the figure. ATL.L and ATL.R represent the left and right anterior temporal lobe, respectively, while FFA.L and FFA.R represent the left and right fusiform face area, respectively. The right side of the figure shows the beta value curves for the left/right ATL (a) and FFA (b) in the pre-surprise and post-surprise conditions of the attribute amnesia task, illustrating the changes in BOLD signal activation related to face perception while it is attended but either without or with memory. Error bars represent SEM. * *p* < 0.05, \*\**p* < 0.01.

To investigate neural responses to face perception, we performed an ROI analysis to calculate beta values for the two conditions. In each ROI, the time courses of beta values for each condition are displayed on the right side of Fig. 3. In the left ATL, two beta values (at 6s and 8s) in the pre-surprise condition were lower than those in the post-surprise condition. A similar pattern was observed in the right ATL, where three beta values (at 6s, 8s, and 10s) in the pre-surprise condition were lower than those in the post-surprise condition. These beta value curves indicates that, compared with the post-surprise condition, bilateral ATL exhibits lower peak activation and earlier decay in the pre-surprise condition. However, no differences in activation pattern were observed in the left and right FFA between the two conditions. The lower neural responses in the ATL during the pre-surprise condition suggest that identity representations might be inhibited, whereas category representations might not be.

### SMG inhibits the activation of ATL in the pre-surprise condition

To investigate the neural circuits underlying the dissociation between attention and WM—particularly how the brain suppresses attended representations that are unnecessary for memory—we applied dynamic causal modeling (DCM) analysis^25^ to examine effective connectivity among key brain regions. This approach enables us to validate the hypothesis that the SMG inhibits face perception to prevent its entry into working memory, while also considering an alternative argument suggesting that all attended information is initially encoded into working memory and subsequently removed if unnecessary from memory storage^3,26–28^. The DCM analysis focused on connectivity changes among three key regions identified in prior analyses: the inhibition region (SMG), the face perception region (ATL or FFA), and the WM storage region (dlPFC)^29^.

In the first stage of the DCM analysis (Fig. 4a), we defined 15 different models among three regions to model the modulatory effect of the pre-surprise condition using the post-surprise condition as the baseline, and then fitted each model for each participant^30^. In the second stage, we compared 15 different models at the group level by calculating the exceedance probability of each model^31^. We then selected the best model using random effects Bayesian Model Selection^32^. The results demonstrated that for both the ATL or FFA, the 15th model best explained the modulatory effect of the pre-surprise condition.

**Fig. 4.**
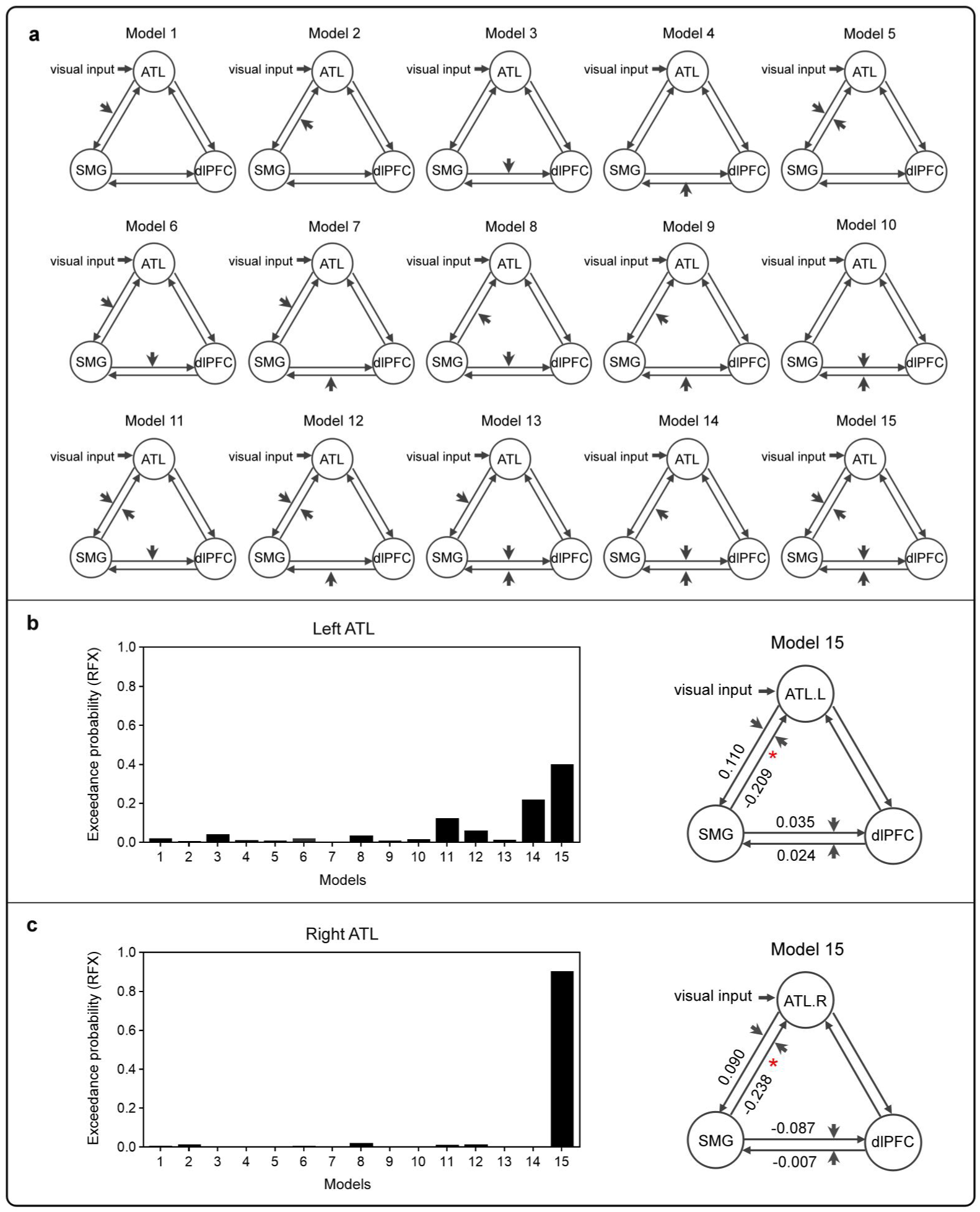
DCM results for inhibitory circuits regulating face identity representation. (**a**). Fifteen different models among the ATL, SMG, and dlPFC were defined for modeling the modulatory effect of the pre-surprise condition on face identity processing. The short gray arrows illustrate the modulatory effect of the pre-surprise condition on interregional connectivity, including the connections from the ATL to the SMG, from the SMG to the ATL, from the SMG to the dlPFC, and from the dlPFC to the SMG. (**b**). The left side of the figure illustrates the exceedance probabilities of the 15 models between SMG, left ATL and dlPFC. The right side illustrates the strength of the modulatory effect on interregional connectivity in the selected Model 15 for the left ATL and its significance level. * *p* < 0.05. (**c**). Similar to Fig.b, but here ATL.R represents the right ATL.

Specifically, for the model of ATL, the pre-surprise condition significant increased the suppression from the SMG to the left ATL (mean values of 49 participants = -0.209, *t*(48) = - 2.462, *p* = 0.017, one-sample t-test, Fig. 4b), as well as to the right ATL (mean values of 49 participants = -0.238, *t*(48) = -2.199, *p* = 0.033, one-sample t-test, Fig. 4c). No significant modulatory effect was found for the other connections in the selected 15th model. As Fig. 4 shows, the brain models indicate a significant inhibitory effect from the SMG to the ATL under the pre-surprise condition. Additionally, for the model of FFA, there was no significant modulatory effect on either the connectivity from the SMG to the left/right FFA or any other inter-regional connectivity (Supplementary Fig. 1). This aligns with the ROI analysis for face perception activation patterns, providing further evidence that neural representations of face identity are inhibited in the pre-surprise condition, while category representations remain unaffected.

Collectively, these findings demonstrate that the SMG functions as a key hub underlying the dissociation mechanism, selectively suppressing the perceptual encoding of unnecessary attentional representations, such as face identity, and preventing their entry into working memory. These results support our hypothesis that such representations are inhibited and blocked from the memory system, rather than the alternative argument that they are initially encoded into the memory system and subsequently suppressed from storage.

### Anodal tDCS on the SMG strengthens inhibitory control over attended inputs

To provide causal validation for our previous neural evidence that attention and WM encoding are dissociable, we applied transcranial direct current stimulation (tDCS) to modulate the SMG, which is the key region enabling this dissociation, aiming to alter its inhibitory control over attended-without-memory information (fully attended but without the necessity for memory report) and assess the resulting changes in its access to WM encoding.

To effectively evaluate changes in its access to WM encoding, we used a modified change-detection paradigm, where stronger inhibitory control results in weaker memory traces of attended information, which can be sensitively measured through reaction time changes^12^. As a basis for understanding our approach, in the classic design of the change-detection paradigm, a memory item and a test item are sequentially presented, and participants are asked to report whether one feature, such as color (i.e., the memory feature), in the test item has changed compared with the memory item. Although changes in another feature, such as identity (i.e., the irrelevant feature), are ignored, reaction times (RT) tend to be faster when both items share the same identity^12,33^. This suggests that the memory trace of the identity feature automatically affects the processing of the subsequent item^34^. Building on this, a modified version of the paradigm introduced a preliminary question about the identity of the memory item^12^. Only after confirming the identity did participants proceed to the color detection task. In this case, the identity is the attended-without-memory feature. The results showed no significant RT difference between same-identity and change-identity conditions, suggesting that attended-without-memory information had been actively inhibited, leading to an even weaker memory trace than ignored information in the classic design.

In the first tDCS experiment of the current study, we utilized a modified change-detection paradigm with colored faces as stimuli, where participants performed a color detection task, and the identity of the face served as attended-without-memory information (Fig. 5c). Prior to the task, high-definition tDCS (Soterix Medical, New York, USA) was applied to target the inhibitory control region, the SMG, identified from the fMRI task. The central electrode was positioned over CP6, delivering 2 mA of current for 20 minutes^35^ (Fig. 5a, Fig. 5b ; see Methods for more details). Ninety participants were divided into three groups: anodal tDCS (enhancing SMG activation), cathodal tDCS (reducing SMG activation), and sham tDCS (as control)^36^. The modulatory effects of tDCS were assessed by comparing active stimulation to sham stimulation^37^. As shown in Fig. 5d, we assessed the effect of anodal stimulation by comparing the anodal group with the sham group and observed a significant interaction (F_1,58_ = 6.465, *p* = 0.014, ŋp^2^ = 0.100). The simple effects test showed that within the anodal group, RT for the same-identity condition was significantly longer than for the change-identity condition (mean difference = 22.21 ms, F_1,58_ = 9.379, *p* = 0.003, ŋp^2^ = 0.139), indicating a stronger inhibitory effect on the attended face identity and a reduction in memory traces. Conversely, in the sham group, there was no significant difference in RT between the two conditions (mean difference = 3.87 ms, F_1,58_ = 0.284, *p* = 0.596, ŋp^2^ = 0.005), reflecting a baseline level of memory traces of the identity^12^. However, no significant interaction was found between the cathodal and sham groups (F_1,58_ = 0.254, *p* = 0.616, ŋp^2^ = 0.004), suggesting no discernible differences in the inhibitory effect following cathodal stimulation.

**Fig. 5.**
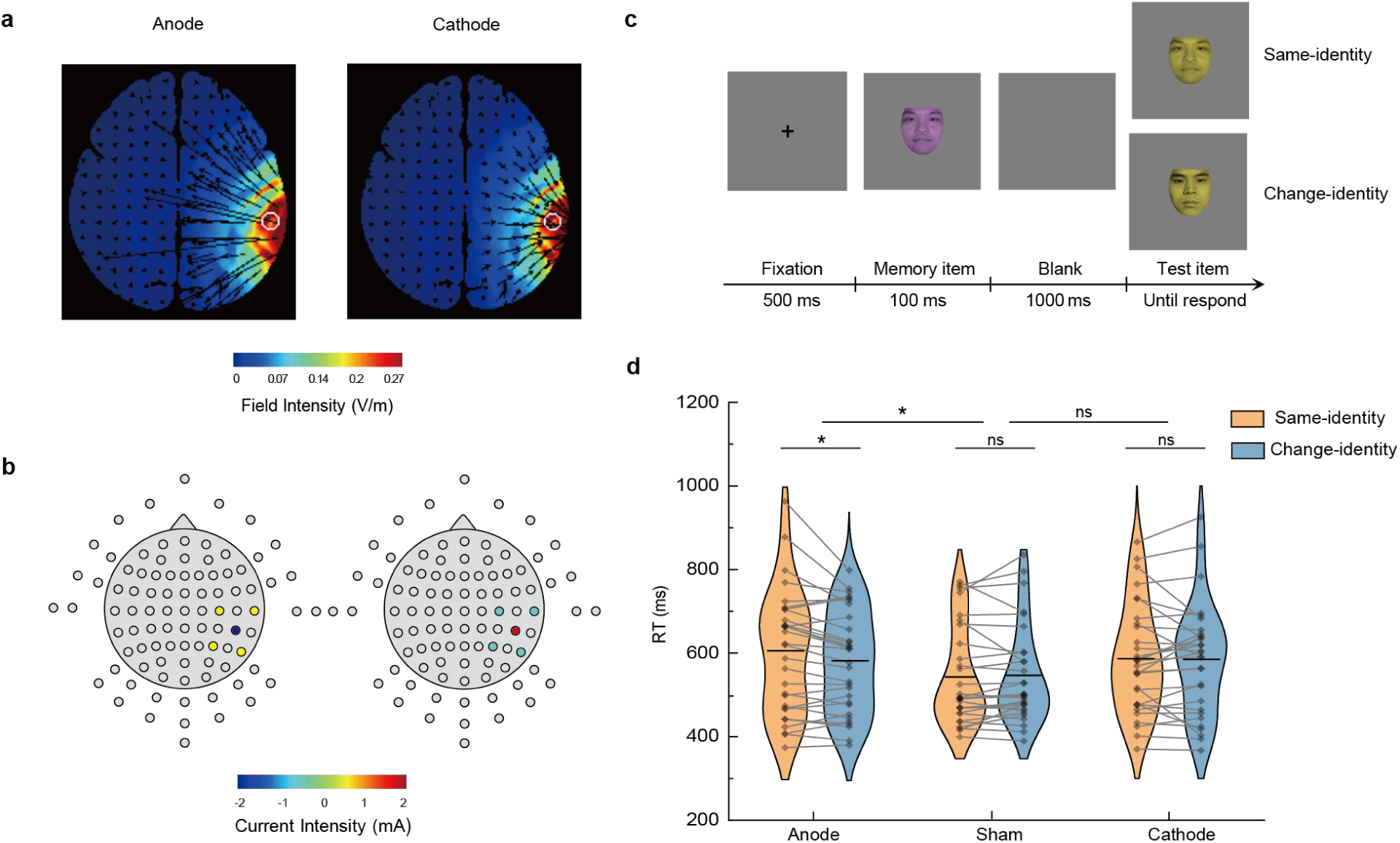
tDCS modulation setup and results of the first tDCS experiment. (**a**) The direction and intensity of the electrical currents were optimized for high-resolution anodal and cathodal stimulation targeting the SMG. (**b**) Electrode configuration, showing the central electrode at CP6, surrounded by electrodes at C4, P4, P8, and T8, along with the applied current intensity. (**c**) Procedure of the modified change-detection task. Participants first judged whether the memory item’s identity matched a pre-specified one. If it did not, they had to memorize its color and report whether the color of the test item differed from that of the memory item. The identity of the memory item served as attended-without-memory information. (**d**) Change-detection task results across three post-tDCS groups. RT (reaction time): a longer RT in the same-identity condition indicates stronger inhibition of attended face identity. Violin plots show mean RT (horizontal line) and individual RTs (scatter points) linked across conditions for each participant. \**p* < 0.05; ns = not significant.

These results highlight that the inhibitory control over the attended identity was markedly stronger in the anodal group than in the sham group, indicating a significant enhancement of SMG inhibitory function and reducing attended identity access to WM encoding following anodal stimulation.

Moreover, to further validate our findings and extend their generalizability, we expanded our investigation of the dissociation mechanism to include simpler stimuli. In the second tDCS experiment, we replicated the procedure from the first experiment, replacing face stimuli with colored shapes. Ninety new participants performed this modified change-detection task, with the shape identity serving as the attended-without-memory information (Fig. 6a). As shown in Fig. 6b, a significant interaction was found between the anodal and sham groups (F_1,58_ = 4.518, *p* = 0.038, ŋp² = 0.072). The anodal group exhibited a significantly longer RT in the same-shape condition compared to the change-shape condition (mean difference = 15.14 ms, F_1,58_ = 6.004, *p* = 0.017, ŋp² = 0.094), indicating stronger inhibitory effects on the attended shape. In contrast, the sham group showed no significant difference between conditions (mean difference = 3.43 ms, F_1,58_ = 0.309, *p* = 0.581, ŋp ² = 0.005). No significant interaction was observed between cathodal and sham groups (F_1,58_ = 1.039, *p* = 0.312, ŋp² = 0.018). These findings align with the first tDCS experiment, suggesting that anodal tDCS enhances the inhibitory control on attended-without-memory information, providing causal evidence for the dissociation mechanism across different types of stimuli.

**Fig. 6.**
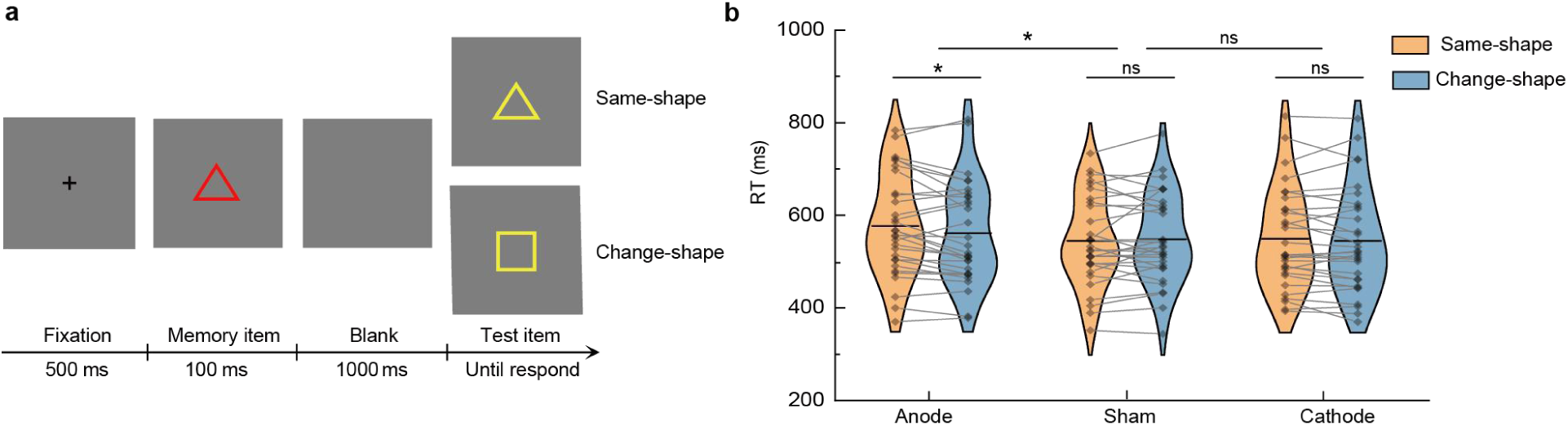
The second tDCS experiment. **(a)** The procedure was identical to the first experiment, but colored shapes replaced colored faces. Participants first judged whether the memory item’s shape was a circle. If it was not, they memorized its color and reported whether the color of the test item had changed relative to the memory item. The shape identity of the memory item served as attended-without-memory information. **(b)** Ninety new participants were divided into three groups, and the results of the change-detection task after tDCS stimulation are shown for each group. \**p* < 0.05; ns = no significance.

## Discussion

This study provides multimodal evidence demonstrating that attentional selection and WM encoding are neurally dissociable processes, and identifies the underlying mechanisms that regulate the transition of attended neural representations into WM. Specifically, neuroimaging evidence combined with dynamic causal modeling identifies the supramarginal gyrus (SMG) as the key region enabling this dissociation, which actively inhibits neural representations of attentional inputs unnecessary for memory, thereby preventing their entry into WM. Furthermore, findings from two tDCS experiments provide causal validation of this dissociation mechanism, showing that enhancing SMG activation strengthens its inhibitory control over attended inputs and reduces their access to WM encoding.

These findings challenge the long-standing view of the relationship between attention and WM encoding, demonstrating that they constitute two dissociable stages of information selection rather than an inseparable process. While previous studies have suggested a shared neural basis for attention and WM encoding^3–6,38^, these results may stem from experimental designs that fail to disentangle these processes. Through an innovative experimental design, we are able to directly investigate the neural dissociation between attentional selection and WM encoding. Crucially, the task design ensured that the target remained at the focus of attention, yet behavioral results showed that this attended information was not encoded into WM when participants had no expectation to report it. This observation, along with prior research on attribute amnesia^11^, provides a solid foundation for establishing the neural separation between these two processes. Building on this, we identify the SMG as the central hub of the dissociation mechanism between attention and WM encoding through its active inhibition of attended inputs that are unnecessary for memory storage. Consistent with our findings, previous neurophysiological and lesion studies provide additional support for the SMG’s role in inhibitory control within cognitive processes^39–43^.

The dissociation mechanism redefines our understanding of how WM encodes attended inputs, demonstrating the presence of a dedicated gating mechanism for WM encoding rather than automatic encoding. The traditional view of WM gating suggests that it automatically encodes attended inputs and later actively suppresses the neural activity of outdated content, facilitating its removal from WM to optimize storage capacity(Gazzaley & Nobre, 2012; Kim et al., 2020; Lewis-Peacock et al., 2018). In contrast, our findings reveal a distinct WM gating mechanism that underlies the regulation of attended neural representations as they transition into WM. Specifically, our dynamic causal modeling results show that the SMG exerts inhibitory control over the ATL, which governs perceptual encoding, rather than the dlPFC, which is associated with memory storage. These findings suggest that, rather than relying on storage-based removal as previously proposed, this gating mechanism selectively inhibits unnecessary perceptual inputs to prevent their entry into WM. Supporting this interpretation, a parallel study demonstrated that attended information without memory requirements minimally impacts memory capacity in a dual-task design and lacks an online memory trace in electroencephalography^44^. Additionally, diffusion tensor imaging tractography identified anatomical pathways supporting inhibitory signal transmission from the SMG to the ATL, providing structural evidence for this gating mechanism^45,46^.

Furthermore, the gating mechanism for attentional inputs into WM exhibits neural plasticity, as demonstrated by our tDCS findings. Enhancing SMG activation amplifies its inhibitory control over attended inputs that do not require memory, providing causal evidence for this mechanism. Specifically, anodal tDCS enhances activity in the stimulated region^36^ and strengthens its functional connectivity with other regions^47,48^, likely reinforcing the synaptic transmission of inhibitory signals from the SMG to cortical areas representing non-memorized attentional inputs. As shown in Fig. 5d, sham tDCS maintained balanced inhibitory control, keeping the neural representations of attended identities at baseline levels, consistent with previous behavioral findings^12^. In contrast, anodal tDCS heightened suppression of these attended representations, reducing them below baseline, leading to weaker memory traces for attended face identities than for unattended ones. A follow-up tDCS experiment confirmed this effect, showing a consistent amplification of inhibitory gating over shape identities (Fig. 6), demonstrating the generalizability of this mechanism from complex to simpler stimuli. These findings underscore the importance of a balanced gating mechanism, as excessive suppression may not benefit memory efficiency. Importantly, the observed functional plasticity highlights the potential for targeted interventions to regulate attentional inputs into WM, offering insights into addressing memory deficits caused by insufficient suppression of irrelevant inputs.

Additionally, the brain’s gating of attended inputs that do not require WM storage is selective. According to our findings, when face reporting is not required, this gating mechanism selectively suppresses identity representations in the ATL, whereas category representations in the FFA remain unaffected. This selectivity raises a critical question: why doesn’t the brain inhibit all attended representations unnecessary for memory to further reduce interference with WM? Previous research suggests that inhibition is an active suppression process that requires substantial executive control resources^49–51^, with inhibitory circuits mediating synaptic transmission of neurotransmitters like GABA demanding significant energy expenditure^45,52,53^. Thus, the gating mechanism likely incurs considerable energy costs. At the same time, encoding unnecessary information into WM can create interference and compete for limited processing capacity. To optimize resource utilization, the brain appears to prioritize the inhibition of high-interference information, such as identity representations that occupy substantial capacity, while preserving lower-interference information, such as basic-level categories, which exert minimal cognitive load^13,54^. This selective inhibition reflects an adaptive trade-off between minimizing interference and conserving cognitive resources.

One issue warrants mention. Anodal tDCS demonstrated a significant enhancement effect; however, the expected reduction effect from cathodal tDCS was not observed, aligning with polarity-specific responses reported in prior studies^55–57^. The absence of a significant cathodal effect could be due to compensatory mechanisms within brain networks^56,58^, or it could result from our use of an offline approach involving only a single session of electrical stimulation prior to the behavioral test^59,60^. Future studies could employ an online approach where stimulation is delivered concurrently with behavioral testing, or employ multiple stimulation sessions to amplify the modulation effect.

## Methods

### Participants

All participants had normal or corrected-to-normal visual acuity, no deficits in facial recognition or color vision, and had not previously participated in similar experiments. The study was approved by the Research Ethics Board of the School of Psychology at South China Normal University. All participants signed informed consent forms prior to participation in the study. A total of 55 participants were recruited for the fMRI study. Six participants were excluded from the analysis, including three who did not complete fMRI scanning and three who exhibited excessive head motion (i.e., over 3 mm in translation or 3 degrees in rotation). The remaining 49 participants (31 females and 18 males, mean age: 20.18 ± 2.01 years) were included in fMRI analyses. In the first tDCS experiment, 98 participants were initially recruited. Eight were excluded either for requesting to abort the experiment or for having a mean RT that exceeded 2.5 SD from the group mean. This left 30 participants in the anodal group (14 females and 16 males, mean age 20.53 ± 1.61 years), 30 in the cathodal group (12 females and 18 males, mean age 20.53 ± 1.36 years), and 30 in the sham group (19 females and 11 males, mean age 20.63 ± 1.79 years). Similarly, in the second tDCS experiment, 97 participants were recruited. Seven were excluded using the same criteria. The remaining participants were divided into an anodal group (30 participants: 19 females and 11 males, mean age 20.23 ± 1.76 years), a cathodal group (30 participants: 22 females and 8 males, mean age 20.47 ± 2.22 years), and a sham group (30 participants: 19 females and 11 males, mean age 20.50 ± 2.27 years).

### Attribute amnesia task in fMRI experiment

The modified attribute amnesia task was conducted using a slow event-related design, comprising four runs (as shown in Fig.1a). Each run consisted of 32 trials, with an inter-trial interval of was 8-12 s. As shown in Fig.1b, each trial began with the fixation display, which consisted of a black central fixation cross and four placeholder circles positioned in the four corners of an invisible square for 1 s. Subsequently, four images including one face and three houses appeared for 1 s, which were replaced by the masks for 100 ms. A blank screen with a fixation cross followed, which presented for varying durations between 8-12 s. The response tasks differed in regular trials and surprise trials. In the regular trial, participants were instructed to report the location of the target face by pressing one of four number keys (1-4) corresponding to the four locations, then the feedback shown for 500 ms. In the surprise trial, the instruction for the face selection task was presented for 2 s to prompt participants to read it before responding, then below the instruction, four different faces appeared randomly with the numbers 1-4 until participants reported the target identity by pressing a corresponding number key. After the face selection task, participants were also asked to report the target location. Subsequently, the feedback for both tasks was displayed for 800 ms. The stimuli contained a total of four stranger-face pictures and 12 house pictures, which were processed into the same shape and grayscale^61^. The external features of faces (including hair or natural overall contours) have been removed, with the internal features of images (including eyes, nose, and mouth) serving as the primary source of information for the perceptual processing of faces^15,62^. The copyright for the Chinese Facial Affective Picture System used in the study has been duly purchased^61^.

### Face-localizer task in fMRI experiment

After the attribute amnesia task, a face-localizer task was performed to localize the brain regions activated during face perception^24^. The localizer task comprised four block-designed runs, each consisting of ten blocks (five face blocks and five house blocks) in randomized order. blocks were separated by 12 s rest periods during which a fixation cross was presented. Each 12 s block comprised 24 trials, with a picture presented for 400 ms followed by a fixation cross for 100 ms. Participants were asked to perform a one-back identification task, judging whether the current picture was the same as the previous one, to maintain their focus on the stimuli.

### fMRI data acquisition

The MRI data were acquired using a Siemens 3T Magnetom Prisma Fit scanner. Blood oxygen level-dependent (BOLD) signals were measured with an echo-planar imaging (EPI) sequence (TR/TE/flip angle = 2,000 ms / 30 ms / 90°, FOV = 192 × 192 mm^2^, matrix size = 64 × 64, slice thickness = 3 mm, number of slices = 32). A high-resolution T1-weighted structural image (TR/TE/flip angle = 2,530 ms / 2.27 ms /7°, FOV = 256 × 256 mm^2^, matrix size = 256 × 256, slice thickness = 1 mm, number of slices=208) was acquired for each subject. The stimuli for the EPI scans were back-projected via a video projector (refresh rate: 60 Hz; spatial resolution: 1,024 × 768) onto a translucent screen placed inside the scanner bore using MATLAB 2018b with Psychtoolbox-3^63^. The participants viewed the stimuli through a mirror located above their eyes. Each participant completed four runs of the attribute amnesia task and four runs of the face-localizer task.

### fMRI data preprocessing

The AFNI software package (version 21.2.03, http://afni.nimh.nih.gov/afni) was employed for fMRI data analysis. The fMRI preprocessing steps included: slice timing correction; head motion correction using realignment on functional volumes; co-registration of the functional image with the structural image; non-linear transformation to the Montreal Neurological Institute (MNI) template; functional volumes were resampled into 3 × 3 × 3mm^3^ resolution; volumes with excessive motion were censored (marked as zeros, otherwise as ones) if the Euclidean norm of derivatives of motion parameters exceeded 0.3 mm; spatial smoothing was applied using a Gaussian filter with a 4 mm full-width half-maximum; and finally, the voxel time series were scaled to percent signal change.

### Whole-brain analyses in the attribute amnesia experiment

A whole-brain analysis was conducted using a general linear model (GLM) with the TENT function of 3dDeconvolve in AFNI^64^. The GLM analyzed the regressors of two experimental conditions (pre-surprise condition: the stimulus onset times of the pre-surprise regular trials, and the post-surprise condition: the stimulus onset times of the post-surprise regular trials) to estimate the event-related BOLD signals after stimulus onset. To control for the baseline and constrain the fitting, three additional task regressors (response onset times of pre-surprise regular trials, response onset times of post-surprise regular trials, as well as the stimuli and response onset times of all surprise trials), six motion regressors (three rotations and three translations), and one binarized censored time series (0 indicates censored time points, and 1 indicates non-censored) were included, resulting in a GLM with 12 regressors.

All regressors were convolved with the canonical hemodynamic response function (HRF) to estimate the BOLD signals for each event. The TENT function, a piecewise linear spline function, was used to flexibly estimate the event-related BOLD time courses. The time courses with seven time points (from 0 to 12s with the time step equal to TR 2s) were modeled by TENT (0,12,7) for each event, and seven beta values corresponding to the time points were obtained. The signal value for the two conditions was defined using the average of the beta value of the peak and the preceding and following time points (4s, 6s, and 8s in the time course)^65,66^. For group analysis, voxel-wise paired t-tests were performed, and a whole-brain t-test map was obtained for the contrast “pre-surprise condition > post-surprise condition” (*p* < 0.01, FDR correction at voxel level, cluster size > 40 voxels).

### Localization and activation analysis of face perception regions

To localize the ATL and FFA, we employed GLM analysis to contrast the activation to face versus house stimuli in the face-localizer experiment. The GLM with the BLOCK (12,1) function of 3dDeconvolve was used to estimate BOLD signal changes from baseline for each event condition. The GLM analysis comprised nine regressors, including two task regressors (onset times of face blocks and house blocks), six motion regressors (three rotations, three translations), and one binarized censored time series (zero indicated censored time points, and one indicated non-censored). A paired t-test map at the group level was obtained for the contrast “face > house” (threshold *p* < 0.01, FDR correction at the voxel level). According to previous studies, the localization criterion for the ATL involved the activated voxel cluster in the anterior part of the temporal lobe, while for the FFA, it was the activated voxel cluster in the middle fusiform gyrus^20^ (Collins & Olson, 2014). Following these criteria, two ROIs (Fig. 3) were extracted from the group level GLM results: bilateral ATLs are responsible for processing face identity feature^20,21^, while bilateral FFAs are involved in processing categorical face information^22,23^.

To further investigate the neural responses to face perception in the pre-surprise and post-surprise conditions, we performed a ROI analysis^24^. We used left/right ATL and FFA separately as masks to calculate the beta values of the two conditions based on the GLM results of the attribute amnesia experiment^65^. For each ROI, the mean beta values at each TR shown in Fig.3.

### Dynamic causal modeling analysis

Prior to the DCM analysis, three key regions were identified as ROIs: the SMG (active inhibition region), the left/right ATL or FFA (face processing regions) and the dlPFC (the most representative working memory processing region)^29^. The mean time series for each ROI was extracted from our preprocessed functional data for the subsequent analyses.

A DCM analysis of the effective connectivity between ROIs under the modulatory effects of the pre-surprise condition was conducted using SPM12 toolbox^25^. The analysis was performed in two stages. In the first stage, the model estimation process involved identifying the parameters that achieved an optimal balance between accuracy and complexity^30^. DCM analysis involves three sets of parameters: parameters C represent the driving input, which refers to the stimuli input into one or more regions, causing change in neural activity throughout the network; parameters A denote the baseline intrinsic connections between the regions influenced by the driving input; parameters B indicate the modulatory effects of experimental manipulations on the specified intrinsic connections. In the second stage, Bayesian model comparison was applied to identify the model that best explained the data, corresponding to the model with the minimal discrepancy between predicted and observed time series.

In our DCM analysis, as shown in Fig. 4a, the first stage involved considering parameters C, A, and B. For parameters C, we investigated the visual influence of the pre-surprise and post-surprise conditions on neural activity within the connectivity network, setting the stimuli onsets of both conditions as driving input to the left/right ATL. For parameters A, we used full connections, including inter-region and self-connections of regions^30,67^. For parameters B, we set the pre-surprise condition as modulatory input to assess its effect on the connectivity between SMG and the other two ROIs^68^. Specifically, based on our hypotheses, we focused on the modulatory effect of the pre-surprise condition on four directional connections: the bidirectional connection (i.e., feedforward and feedback) between SMG and ATL, as well as between SMG and dlPFC. The modulatory input could affect the connection from ATL to SMG (model 1); from SMG to ATL (model 2); from SMG to dlPFC (model 3); from dlPFC to SMG (model 4); two of the four connections (models 5-10); three of the four connections (models 11-14); and all four connections (model 15).

In the second stage, model comparison was performed by calculating the exceedance probability of each model and using the Random Effects Bayesian Model Selection^31,32^. This process evaluated the probability that a given model was more likely than any other included model from the generated data from randomly selected participants. Following the selection of the optimal model, we extracted all the modulatory effect values of the selected model from 49 individuals and analyzed the modulatory effect on each inter-regional connection using one-sample t-test.

The DCM analysis for face category processing followed a similar procedure to that for face identity processing, except that the ATL was replaced by the FFA, as shown in Supplementary Fig. 1.

### HD-tDCS data acquisition

High-definition transcranial direct current stimulation (HD-tDCS), as an advanced, non-invasive brain stimulation technique, modulates cortical excitability in a focused and intensive manner by concentrating the direct current within a defined ring over the target areas^69^. Typically, anodal tDCS enhances cortical excitability, cathodal tDCS reduces it, and sham tDCS serves as the control^36^. In our study, we utilized HD-tDCS (Soterix Medical, New York, USA) to modulate the excitability of the SMG (coordinates as illustrated in Supplementary Table 1). The electrode montage was designed using HD-Explore Version 6.0 software with the MNI-152 template to optimize current flow to the SMG and estimate the induced electric field (Fig. 5a). For anodal tDCS, a 4 × 1 multielectrode anode-center cathode-surround montage was used, centered on the right SMG (CP6), with surrounding electrodes at C4, P4, P8, and T8, following the 10-10 electrode site placement^35^. Conversely, a cathode-center anode-surround montage was used for cathodal tDCS. Participants in the active group underwent 20 minutes of 2.0 mA stimulation with a 30 s ramp up and down). In the sham group, the electrodes were designed with half as in the anodal group and half as in the cathode group, but no stimulation was applied for 20 minutes except for a 30 s ramp up and a 30 s ramp down at the beginning and end of the procedure.

Following tDCS stimulation, participants performed the change-detection task and then completed the Questionnaire of Sensations Related to Transcranial Electrical Stimulation^70^. This questionnaire recorded the sensations experienced by participants during the tDCS session, such as itching, pain, and burning. In the first tDCS experiment, the questionnaire analysis revealed no significant differences in reported sensations between the Anode, Sham, and Cathode groups (F_2,87_ = 2.364, *p* = 0.104). Similarly, in the second experiment, no significant differences were found between the tDCS groups (F_2,87_ = 1.214, *p* = 0.302). These results suggest that the tDCS setup successfully blinded participants, preventing them from distinguishing between active and sham stimulation^37^.

### Change-detection task

We employed the modified change-detection paradigm^12^ to evaluate the impact of tDCS modulation of the SMG on the inhibition of attended-without-memory information. During the experiment, participants were seated approximately 50 cm from a 17-inch CRT monitor (60 Hz, 1024 × 768 screen resolution) and responded using a computer keyboard. The display background was medium gray (RGB: 128, 128, 128). In the first tDCS experiment, all stimuli in the memory and test displays were colored faces. There were five faces of each gender, each with five different colors: blue (91, 115, 213), green (112, 173, 71), yellow (255, 192, 0), red (192, 0, 0), and purple (112, 48, 160), all presented with 40% transparency. In the second tDCS experiment, the stimuli were colored shapes, which included circles, hexagons, triangles, squares, and stars. The colors were randomly selected from blue (0, 0, 255), green (0, 128, 0), yellow (255, 255, 0), red (255, 0, 0) and pink (255, 192, 203).

In the first tDCS experiment, participants completed a total of 120 trials, randomized into three blocks. In each trial (Fig. 5c), after a 500 ms fixation display, a memory item with a randomly selected colored face was presented in the center of the screen for 100 ms. Following a 1000 ms blank interval, a test item with a colored face was presented at the center of the screen until the participant responded or 2000 ms elapsed. Participants first judged whether the memory item’ s identity matched a pre-specified person. If it did, they only had to report whether the test item was also the same person (catching trials, would be excluded from the analysis). If the identity did not match, participants had to memorize the color of the face and report whether the color of the test item had changed compared to the memory item. In these non-match trials, the identity of the memory item was fully attended to but not required for reporting (i.e., attended-without-memory information). Critically, in half of the trials (same-identity condition), the identity of the test item remained the same as that of the memory item, whereas in the other half of the trials (change-identity condition), the identity of the test item changed. In the second tDCS experiment (Fig. 6a), the procedure was identical to the first, with colored shapes replacing the face stimuli. Participants first judged whether the memory item was a circle, making the shape identity the attended-without-memory information. If the item was a circle, they reported whether the test item was also a circle (catching trials). If it was not, they memorized its color and reported whether the color of the test item was different from that of the memory item.

### Analysis for change-detection task in the tDCS experiment

The identity non-match trials, where participants detected color with identity considered as attended-without-memory information, were analyzed. Catching trials, focused on identity detection, were excluded prior to analysis^12^. In line with previous research, the analysis focused on reaction times, with accuracy above 95%. Trials with incorrect responses and those exceeding 2.5 standard deviations from the mean reaction times for each condition and participant were removed. In the first tDCS experiment, 8.21% of trials were excluded in the anodal group, 7.31% in the cathodal group, and 6.12% in the sham group. In the second experiment, 4.25% of trials were excluded in the anodal group, 5.09% from the cathodal group, and 6.03% in the sham group. Statistical analyses were performed using F-tests, with Greenhouse-Geisser corrections applied when sphericity violations occurred.

## Data availability

The datasets generated and analyzed during the current study are available on https://github.com/LYYpsy/data.

## Code availability

The example code used in this manuscript is available online at https://github.com/LYYpsy/memory-reselection_mechanism/tree/main/code_OSF.

## Acknowledgements

This work was supported by grants from Science and Technology Innovation 2030-“Brain Science and Brain-like Research” Major Project (No.2022ZD0210800 for H.C.), Emerging Enhancement Technology under Perspective of Humanistic Philosophy, supported by National Office for Philosophy and Social Science (No. 20&ZD045 for H.C.), National Natural Science Foundation of China (No.31971032 for P.Q., No.32171046 for H.C., No.32200844 for Y.F.), Fundamental Research Funds for the Central Universities (226-2024-00207, 226-2024-00118).

## Author contributions

Y.L. was responsible for conceptualization, data acquisition, formal analysis, drafting the manuscript, and reviewing and editing. F.Y. and E.T. contributed to conceptualization, validation, drafting the manuscript, and reviewing and editing. H.W., J.H., M.X., and Y.Z. were responsible for data analysis, drafting the manuscript, and reviewing its content. J.H. also contributed to data acquisition. H.L. provided methodology guidance, supervision for analysis, and manuscript review. H.C. and P.Q. were responsible for conceptualization, funding acquisition, methodology, supervision, and manuscript review and editing.

## References

1. Baddeley, A. D. Working memory: theories, models, and controversies. Annu. Rev. Psychol. 63, 1–29 (2012).

2. Panichello, M. F. & Buschman, T. J. Shared mechanisms underlie the control of working memory and attention. Nature 592, 601–605 (2021).

3. Gazzaley, A. & Nobre, A. C. Top-down modulation: bridging selective attention and working memory. Trends Cogn. Sci. 16, 129–135 (2012).

4. Jonikaitis, D. & Moore, T. The interdependence of attention, working memory and gaze control: behavior and neural circuitry. Curr. Opin. Psychol. 29, 126–134 (2019).

5. Zanto, T. P. Causal role of the prefrontal cortex in top-down modulation of visual processing and working memory. Nat. Neurosci. 14, 8 (2011).

6. Majerus, S., Péters, F., Bouffier, M., Cowan, N. & Phillips, C. The Dorsal Attention Network Reflects Both Encoding Load and Top–down Control during Working Memory. J. Cogn. Neurosci. 30, 144–159 (2018).

7. Mendoza-Halliday, D., Xu, H., Azevedo, F. A. C. & Desimone, R. Dissociable neuronal substrates of visual feature attention and working memory. Neuron 112, 850–863.e6 (2024).

8. Awh, E., Vogel, E. K. & Oh, S.-H. Interactions between attention and working memory. Neuroscience 139, 201–208 (2006).

9. Hamblin-Frohman, Z. & Becker, S. I. Attending object features interferes with visual working memory regardless of eye-movements. J. Exp. Psychol. Hum. Percept. Perform. 45, 1049–1061 (2019).

10. Tay, D. & McDonald, J. J. Attentional enhancement predicts individual differences in visual working memory under go/no-go search conditions. PLOS Biol. 20, e3001917 (2022).

11. Chen, H. & Wyble, B. Amnesia for Object Attributes: Failure to Report Attended Information That Had Just Reached Conscious Awareness. Psychol. Sci. 26, 203–210 (2015).

12. Fu, Y., Zhou, Y., Zhou, J., Shen, M. & Chen, H. More attention with less working memory: The active inhibition of attended but outdated information. Sci. Adv. 7, eabj4985 (2021).

13. Rossion, B. & Jacques, C. The N170: Understanding the Time Course of Face Perception in the Human Brain. (Oxford University Press, 2011).

14. Rossion, B. Humans Are Visual Experts at Unfamiliar Face Recognition. Trends Cogn. Sci. 22, 471–472 (2018).

15. Tanaka, J. W. The entry point of face recognition: Evidence for face expertise. J. Exp. Psychol. Gen. 130, 534–543 (2001).

16. Golby, A. J., Gabrieli, J. D. E., Chiao, J. Y. & Eberhardt, J. L. Differential responses in the fusiform region to same-race and other-race faces. Nat. Neurosci. 4, 845–850 (2001).

17. Liu, J., Harris, A. & Kanwisher, N. Stages of processing in face perception: an MEG study. Nat. Neurosci. 5, 910–916 (2002).

18. Tam, J., Mugno, M. K., O’Donnell, R. E. & Wyble, B. And like that, they were gone: A failure to remember recently attended unique faces. Psychon. Bull. Rev. 28, 2027–2034 (2021).

19. Chen, H. et al. Does attribute amnesia occur with the presentation of complex, meaningful stimuli? The answer is, “it depends”. Mem. Cognit. 47, 1133–1144 (2019).

20. Collins, J. A. & Olson, I. R. Beyond the FFA: The role of the ventral anterior temporal lobes in face processing. Neuropsychologia 61, 65–79 (2014).

21. Rajimehr, R., Young, J. C. & Tootell, R. B. H. An anterior temporal face patch in human cortex, predicted by macaque maps. Proc. Natl. Acad. Sci. 106, 1995–2000 (2009).

22. Haxby, J. V. et al. Distributed and Overlapping Representations of Faces and Objects in Ventral Temporal Cortex. Science 293, 2425–2430 (2001).

23. Liu, J., Harris, A. & Kanwisher, N. Perception of Face Parts and Face Configurations: An fMRI Study. J. Cogn. Neurosci. 22, 203–211 (2010).

24. Guo, B. & Meng, M. The encoding of category-specific versus nonspecific information in human inferior temporal cortex. NeuroImage 116, 240–247 (2015).

25. Friston, K. J., Harrison, L. & Penny, W. Dynamic causal modelling. NeuroImage 19, 1273–1302 (2003).

26. Lewis-Peacock, J. A., Kessler, Y. & Oberauer, K. The removal of information from working memory. Ann. N. Y. Acad. Sci. 1424, 33–44 (2018).

27. Kim, H., Smolker, H. R., Smith, L. L., Banich, M. T. & Lewis-Peacock, J. A. Changes to information in working memory depend on distinct removal operations. Nat. Commun. 11, 6239 (2020).

28. Schmicker, M., Schwefel, M., Vellage, A.-K. & Müller, N. G. Training of Attentional Filtering, but Not of Memory Storage, Enhances Working Memory Efficiency by Strengthening the Neuronal Gatekeeper Network. J. Cogn. Neurosci. 28, 636–642 (2016).

29. Owen, A. M., McMillan, K. M., Laird, A. R. & Bullmore, E. N-back working memory paradigm: A meta-analysis of normative functional neuroimaging studies. Hum. Brain Mapp. 25, 46–59 (2005).

30. Zeidman, P. et al. A guide to group effective connectivity analysis, part 1: First level analysis with DCM for fMRI. NeuroImage 200, 174–190 (2019).

31. Zeidman, P. et al. A guide to group effective connectivity analysis, part 2: Second level analysis with PEB. NeuroImage 200, 12–25 (2019).

32. Rigoux, L., Stephan, K. E., Friston, K. J. & Daunizeau, J. Bayesian model selection for group studies — Revisited. NeuroImage 84, 971–985 (2014).

33. Shen, M., Tang, N., Wu, F., Shui, R. & Gao, Z. Robust object-based encoding in visual working memory. J. Vis. 13, 1 (2013).

34. Gao, T., Gao, Z., Li, J., Sun, Z. & Shen, M. The perceptual root of object-based storage: An interactive model of perception and visual working memory. J. Exp. Psychol. Hum. Percept. Perform. 37, 1803–1823 (2011).

35. Koessler, L. et al. Automated cortical projection of EEG sensors: Anatomical correlation via the international 10–10 system. NeuroImage 46, 64–72 (2009).

36. Stagg, C. J. & Nitsche, M. A. Physiological Basis of Transcranial Direct Current Stimulation. The Neuroscientist 17, 37–53 (2011).

37. Tu, Y. et al. Perturbing fMRI brain dynamics using transcranial direct current stimulation. NeuroImage 237, 118100 (2021).

38. van Kerkoerle, T., Self, M. W. & Roelfsema, P. R. Layer-specificity in the effects of attention and working memory on activity in primary visual cortex. Nat. Commun. 8, 13804 (2017).

39. Simonyan, K. Inferior parietal cortex as a hub of loss of inhibition and maladaptive plasticity (S39.002). Neurology 88, S39.002 (2017).

40. Wang, J. et al. Functional topography of the right inferior parietal lobule structured by anatomical connectivity profiles: Structural and Functional Topography of the RIPL. Hum. Brain Mapp. 37, 4316–4332 (2016).

41. Talanow, T., Kasparbauer, A.-M., Lippold, J. V., Weber, B. & Ettinger, U. Neural correlates of proactive and reactive inhibition of saccadic eye movements. Brain Imaging Behav. 14, 72–88 (2020).

42. Bunge, S. A., Dudukovic, N. M., Thomason, M. E., Vaidya, C. J. & Gabrieli, J. D. E. Immature Frontal Lobe Contributions to Cognitive Control in Children: Evidence from fMRI. Neuron 33, 301–311 (2002).

43. Padmanabhan, A. et al. Developmental Changes in Brain Function Underlying Inhibitory Control in Autism Spectrum Disorders. Autism Res. Off. J. Int. Soc. Autism Res. 8, 123–135 (2015).

44. Zhu, P., Fu, Y., Wyble, B., Shen, M. & Chen, H. A new aspect of cognitive selectivity: Working memory reselection for attended information. PsyArXiv (2022).

45. Isaacson, J. S. & Scanziani, M. How Inhibition Shapes Cortical Activity. Neuron 72, 231–243 (2011).

46. Catani, M. & Thiebaut de Schotten, M. A diffusion tensor imaging tractography atlas for virtual in vivo dissections. Cortex 44, 1105–1132 (2008).

47. Bachtiar, V., Near, J., Johansen-Berg, H. & Stagg, C. J. Modulation of GABA and resting state functional connectivity by transcranial direct current stimulation. eLife 4, e08789 (2015).

48. Antonenko, D. et al. Microstructural and functional plasticity following repeated brain stimulation during cognitive training in older adults. Nat. Commun. 14, 3184 (2023).

49. Anderson, M. C. & Green, C. Suppressing unwanted memories by executive control. Nature 410, 366–9 (2001).

50. Vogel, E., McCollough, A. & Machizawa, M. Neural measures reveal individual differences in controlling access to working memory. Nature 438, 500–3 (2005).

51. Oberauer, K., Lewandowsky, S., Farrell, S., Jarrold, C. & Greaves, M. Modeling working memory: an interference model of complex span. Psychon. Bull. Rev. 19, 779–819 (2012).

52. Qin, P. et al. Vascular-metabolic and GABAergic Inhibitory Correlates of Neural Variability Modulation. A Combined fMRI and PET Study. Neuroscience 379, 142–151 (2018).

53. Harris, J. J., Jolivet, R. & Attwell, D. Synaptic Energy Use and Supply. Neuron 75, 762–777 (2012).

54. Bays, P. M., Schneegans, S., Ma, W. J. & Brady, T. F. Representation and computation in visual working memory. *Nat*. Hum. Behav. 8, 1016–1034 (2024).

55. Almeida, J. et al. Polarity-specific transcranial direct current stimulation effects on object-selective neural responses in the inferior parietal lobe. Cortex 94, 176–181 (2017).

56. Jacobson, L., Koslowsky, M. & Lavidor, M. tDCS polarity effects in motor and cognitive domains: a meta-analytical review. Exp. Brain Res. 216, 1–10 (2012).

57. Stagg, C. J. et al. Cortical activation changes underlying stimulation-induced behavioural gains in chronic stroke. Brain 135, 276–284 (2012).

58. Müller, D., Habel, U., Brodkin, E. S. & Weidler, C. High-definition transcranial direct current stimulation (HD-tDCS) for the enhancement of working memory - A systematic review and meta-analysis of healthy adults. Brain Stimulat. 15, 1475–1485 (2022).

59. Batsikadze, G., Moliadze, V., Paulus, W., Kuo, M.-F. & Nitsche, M. A. Partially non-linear stimulation intensity-dependent effects of direct current stimulation on motor cortex excitability in humans. J. Physiol. 591, 1987–2000 (2013).

60. Kuo, H.-I. et al. Comparing cortical plasticity induced by conventional and high-definition 4 × 1 ring tDCS: a neurophysiological study. Brain Stimulat. 6, 644–648 (2013).

61. Gong, X., Huang, Y. X., Wang, Y. & Luo, Y. J. Revision of the Chinese facial affective picture system. Chin. Ment. Health J. 40–46 (2011).

62. Palermo, R. & Rhodes, G. Are you always on my mind? A review of how face perception and attention interact. Neuropsychologia 45, 75–92 (2007).

63. Kleiner, M. et al. What’s new in Psychtoolbox-3? Perception 36, 1–16 (2007).

64. McGuire, J. T. & Kable, J. W. Medial prefrontal cortical activity reflects dynamic re-evaluation during voluntary persistence. Nat. Neurosci. 18, 760–766 (2015).

65. Handwerker, D. A., Ollinger, J. M. & D’Esposito, M. Variation of BOLD hemodynamic responses across subjects and brain regions and their effects on statistical analyses. NeuroImage 21, 1639–1651 (2004).

66. Lindquist, M. A. & Wager, T. D. Validity and power in hemodynamic response modeling: A comparison study and a new approach. Hum. Brain Mapp. 28, 764–784 (2007).

67. Wu, H. et al. Decoding subject’s own name in the primary auditory cortex. Hum. Brain Mapp. 44, 1985–1996 (2023).

68. Huang, L. et al. A source for awareness-dependent figure–ground segregation in human prefrontal cortex. Proc. Natl. Acad. Sci. 117, 30836–30847 (2020).

69. Polanía, R., Nitsche, M. A. & Ruff, C. C. Studying and modifying brain function with non-invasive brain stimulation. Nat. Neurosci. 21, 174–187 (2018).

70. Antal, A. et al. Low intensity transcranial electric stimulation: Safety, ethical, legal regulatory and application guidelines. Clin. Neurophysiol. 128, 1774–1809 (2017).

