## Supplementary Table 1 for "Breaking the Link: Neural Dissociation of Attention and Working Memory through Inhibitory Control"

### **Supplementary material**

Supplementary Table 1. MNI Coordinates for all the regions of interest.

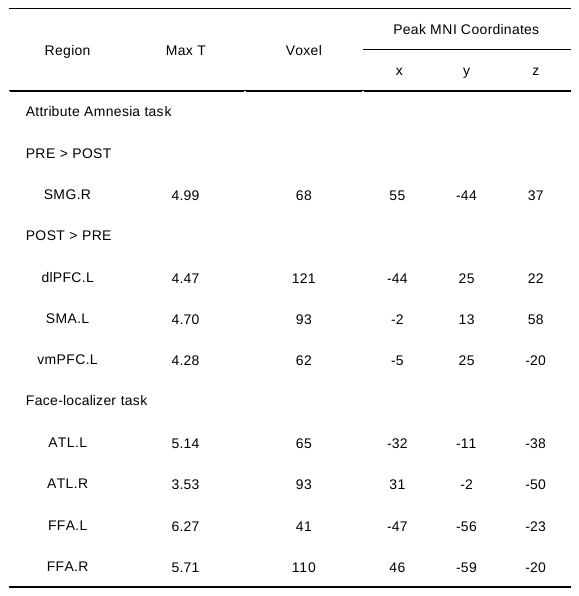

Note: dlPFC.L means left Dorsolateral Prefrontal Cortex, SMA.L means left Supplementary Motor Area, vmPFC.L means left Ventromedial Prefrontal Cortex, SMG.R means right Supramarginal Gyrus, ATL.R means right Anterior Temporal Lobe, FFA.L means left Fusiform Face Area. The coordinate order is LPI.

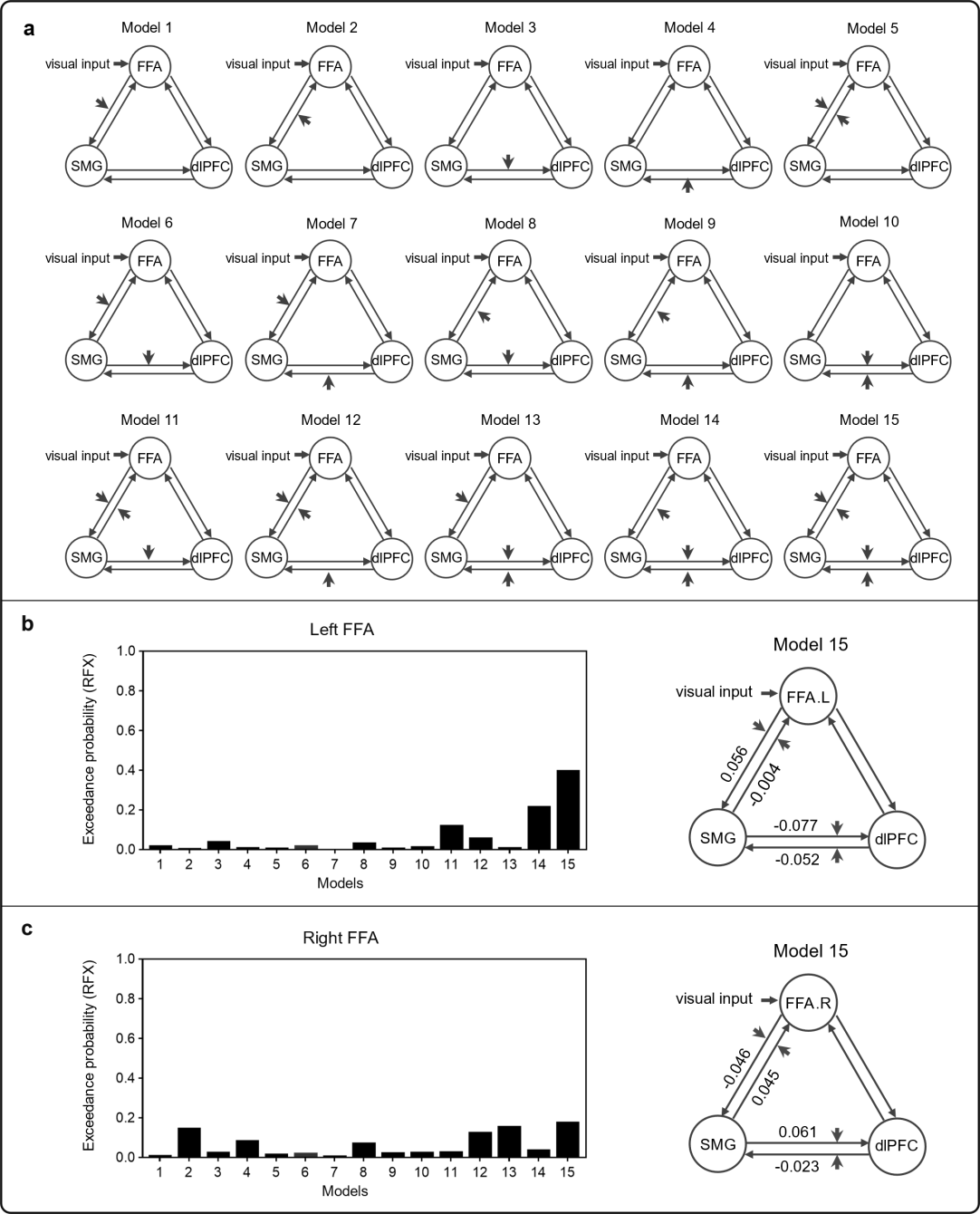

**Supplementary Fig.1. DCM results for inhibitory circuits regulating face category representation.** The analysis followed a similar procedure as in Fig. 4, but the face perception region was replaced by the FFA instead of the ATL.
